# WHEP Domain of Glycyl-tRNA Synthetase Regulates Neuropilin 1 Binding and Vascular Permeability

**DOI:** 10.1101/2025.05.14.654074

**Authors:** Yao Tong, Noemi Gioelli, Huimin Zhang, Calin Dan Dumitru, Sachiko Kanaji, Motoki Horii, Ka Neng Cheong, Jiadong Zhou, Brad Bolon, Ge Bai, Guido Serini, Taisuke Kanaji, Xiang-Lei Yang

**Affiliations:** Department of Molecular Medicine, The Scripps Research Institute, La Jolla, CA, 92037, USA; Candiolo Cancer Institute - Fondazione del Piemonte per l’Oncologia (FPO) Istituto di Ricovero e Cura a Carattere Scientifico (IRCCS), Candiolo (TO), Italy; GEMpath Inc., Longmont, CO 80501, USA; The MOE Frontier Research Center of Brain & Brain-Machine Integration, Zhejiang University School of Brain Science and Brain Medicine, Hangzhou, 310058, China; Department of Oncology, University of Torino School of Medicine, Candiolo (TO), Italy

**Keywords:** glycyl-tRNA synthetase, WHEP domain, neuropilin 1, vascular permeability

## Abstract

Aminoacyl-tRNA synthetases (aaRSs) charge tRNAs with their cognate amino acids, ensuring accurate translation of the genetic code from mRNA to protein. During eukaryotic evolution, aaRSs acquired additional domains with unclear functions, including the WHEP domain, a two-helix bundle found in several eukaryotic aaRSs such as glycyl-tRNA synthetase (GlyRS, encoded by *GARS1*). We generated *Gars1*^ΔWHEP^ mutant mice lacking exon 2, which disrupts most of the WHEP domain. Homozygous *Gars1*^ΔWHEP/ΔWHEP^ mice exhibited late embryonic or neonatal lethality, with delayed lung development, characterized by reduced airway dilation (inflatability), and increased vascular leakage. Disruption of the WHEP domain did not impair tRNA aminoacylation but inhibited the free release of GlyRS from cells. Instead, GlyRS^ΔWHEP^ was found in the membrane fractions and showed a stronger interaction with the extracellular region of neuropilin 1 (Nrp1) receptor compared to full-length GlyRS. This aberrant interaction enhanced Nrp1’s endocytic activity and significantly reduced the localization of the Nrp1 interactor VE-cadherin at the adherens junctions of endothelial cells. A heterozygous knockout of Nrp1 in the *Gars1* ^ΔWHEP/ΔWHEP^ mice partially rescued body weight and vascular permeability defects. This study establishes a physiological role for the GlyRS WHEP domain in lung development and its regulation of GlyRS-Nrp1 interaction and vascular permeability.

## 1. Introduction

Aminoacyl-tRNA synthetases (aaRSs) are a group of enzymes that play a central role in protein synthesis by attaching the correct amino acid to its corresponding transfer RNA (tRNA). This process, known as aminoacylation, is a crucial step in translating the genetic code from mRNA to protein. The 20 different aaRSs, each specific to a particular proteinogenic amino acid, form one of the most ancient protein families and are indispensable for viability of all three domains of life. Two sets of nucleus-encoded aaRSs are necessary for eukaryotes to support mRNA translation that takes place in the cytoplasm and the mitochondria. During the evolution of eukaryotes, cytoplasmic aaRSs have been extended to harbor additional domains that are usually dispensable for tRNA aminoacylation [<u>1</u>], suggesting functions beyond their fundamental involvement in protein synthesis [<u>2</u>].

One such domain is known as the WHEP domain, which forms a two-helix bundle, occurs in six different cytoplasmic tRNA synthetases [<u>3</u>]. (WHEP is an acronym based on the names of four of the six WHEP-containing aaRSs.) In glycyl-, histidyl-, and tryptophanyl-tRNA synthetases (GlyRS, HisRS, and TrpRS), this domain is located at their N-terminus and is appended to the corresponding enzymatic core through a flexible linker [<u>4</u>]. Crystal structure of the human GlyRS/tRNA complex has revealed that the WHEP domain is situated near the single-stranded 3′-end of tRNA, without direct contact [<u>5</u>]. Deletion of the WHEP domain slightly reduced the catalytic efficiency of *Caenorhabditis elegans* and human GlyRS *in vitro*[<u>5</u>], [<u>6</u>]. Nevertheless, the functional impact of the WHEP domain for GlyRS function in animal cells remains unclear.

Many aaRSs have been found in systemic circulation of mammals and have been shown to be released from numerous cell types [<u>7</u>]. Interestingly, through an unbiased interactome study, all three N-terminal WHEP domain-containing aaRSs have also been identified as interaction partners for the receptor neuropilin 1 (Nrp1)[<u>8</u>]. Deletion of the WHEP domain, as in a splice variant of TrpRS (TrpRS^ΔWHEP^, also known as mini-TrpRS)[<u>9</u>], enhances the Nrp1 interaction both in cells and *in vitro*[<u>8</u>]. Several Charcot-Marie-Tooth Type 2D neuropathy (CMT2D)-causing mutation in GlyRS, which could remotely alter the conformation of the WHEP domain relative to the core synthetase [<u>10</u>], have also been demonstrated to enhance the interaction with Nrp1[<u>7b</u>]. Therefore, WHEP domain appears to be a key regulator for the synthetase-Nrp1 interactions.

Nrp1 is a transmembrane protein enriched in neurons and endothelial cells [<u>11</u>]. It is a key regulator for vascular and neuronal development, as Nrp1 knockout mice exhibit severe abnormalities in blood vessels[<u>12</u>] and peripheral nerves[<u>13</u>]. The extracellular region of Nrp1 contains binding sites for both class 3 semaphorins (SEMA3s)[<u>14</u>] and vascular endothelial growth factor A (VEGF-A) [<u>15</u>], through which Nrp1 functions as a co-receptor with type A/D plexin (PLXNA/D) receptors for SEMA3 signaling, and with VEGF receptor-2 (VEGFR-2) for VEGF-A, respectively. Nrp1 also has co-receptor-independent activities in promoting the endocytosis of membrane proteins associated with Nrp1, due to the interaction of its short cytosolic tail with trafficking adaptors, such as GIPC PDZ domain containing family member 1 (GIPC1) [<u>16</u>]. VE-cadherin, a membrane protein located in adherens junctions of endothelial cells, exemplifies this mechanism. Its interaction with Nrp1 facilitates internalization-dependent turnover, which helps to maintain the basal level of vascular permeability [<u>8</u>]. Interestingly, when overexpressed in endothelial cells, TrpRS^ΔWHEP^ acts as a negative regulator for Nrp1-mediated VE-cadherin turnover, which in turn decreases vascular permeability[<u>8</u>].

To probe the physiological role of the WHEP domain in mammals, we generated an exon2-deleted mutant mouse strain in which the majority of the GlyRS WHEP domain was disrupted. These *Gars1*^ΔWHEP/ΔWHEP^ mice exhibit late embryonic or neonatal lethality. At both embryonic day (E) 18.5 and postnatal day (P) 0, the lungs of *Gars1^ΔWHEP/ΔWHEP^* animals exhibit a delayed development phenotype characterized by incompletely dilated airways with pulmonary vascular leakage. The *Gars1^ΔWHEP/ΔWHEP^* embryos also show reduced body weights at E18.5, with some exhibiting anasarca (whole-body edema). No defect in tRNA aminoacylation was found in the lung of *Gars1^ΔWHEP/ΔWHEP^* animals. Interestingly, disruption of the WHEP domain inhibits the free release of GlyRS from cells. Compared with the full-length protein, GlyRS*^ΔWHEP^* has increased association with membrane fractions and interaction with Nrp1. This aberrant interaction promotes the endocytic activity of Nrp1, which controls VE-cadherin turnover and adherens junction stability in endothelial cells. Heterozygous knockout of Nrp1 in the *Gars1*^ΔWHEP^ mice partially restores the body weight and normalizes vascular wall permeability, providing *in vivo* evidence supporting aberrant Nrp1 activation in *Gars1^ΔWHEP/ΔWHEP^* animals. This study suggests a physiological role of GlyRS in regulating blood vessel biology mediated, at least in part, through the Nrp1 receptor and a mechanism that activates Nrp1 via the removal of the appended WHEP domain of GlyRS.

## 2. Results

### 2.1 Generation of WHEP-defective *Gars1* mutant mice

The WHEP domain was first appended to GlyRS at the stage of nematodes and is conserved for the entire animal kingdom (Figure 1a). Among the family members, GlyRS is one of the exceptions where the synthetases responsible for tRNA aminoacylation in the cytoplasm and the mitochondria are encoded by the same gene *Gars1 [*<u>17</u>]. The two forms are differentiated through alternative translation initiation sites encoded within exon 1 (Figure 1b). The mitochondria-targeting GlyRS is translated from an upstream initiation site to incorporate an N-terminal mitochondrial targeting signal (MTS), which is cleaved from the synthetase after its translocation to the mitochondria, resulting in a form identical or similar to the cytosolic GlyRS, although the precise cleavage site of the MTS has not been experimentally established [<u>18</u>]. The cytosolic GlyRS is translated from the second ATG translation initiation site, and the WHEP domain is presented at the N-terminus of the synthetase. The entire WHEP domain spans from exon 1 to exon 3, with exon 2 encoding most of the sequence (i.e., G19-K52) (Figure 1b). Therefore, to disrupt the WHEP domain we deleted exon2 from C57BL/6J mice (Figure 1c). The resulting exon2-deleted allele is designated as *Gars1^Δ2^ or Gars1^ΔWHEP^*, whose expression was demonstrated by Western blot using primary Mouse Embryonic Fibroblasts (MEFs) from both heterozygous and homozygous mutant mice (Figure 1d).

**Figure 1.**
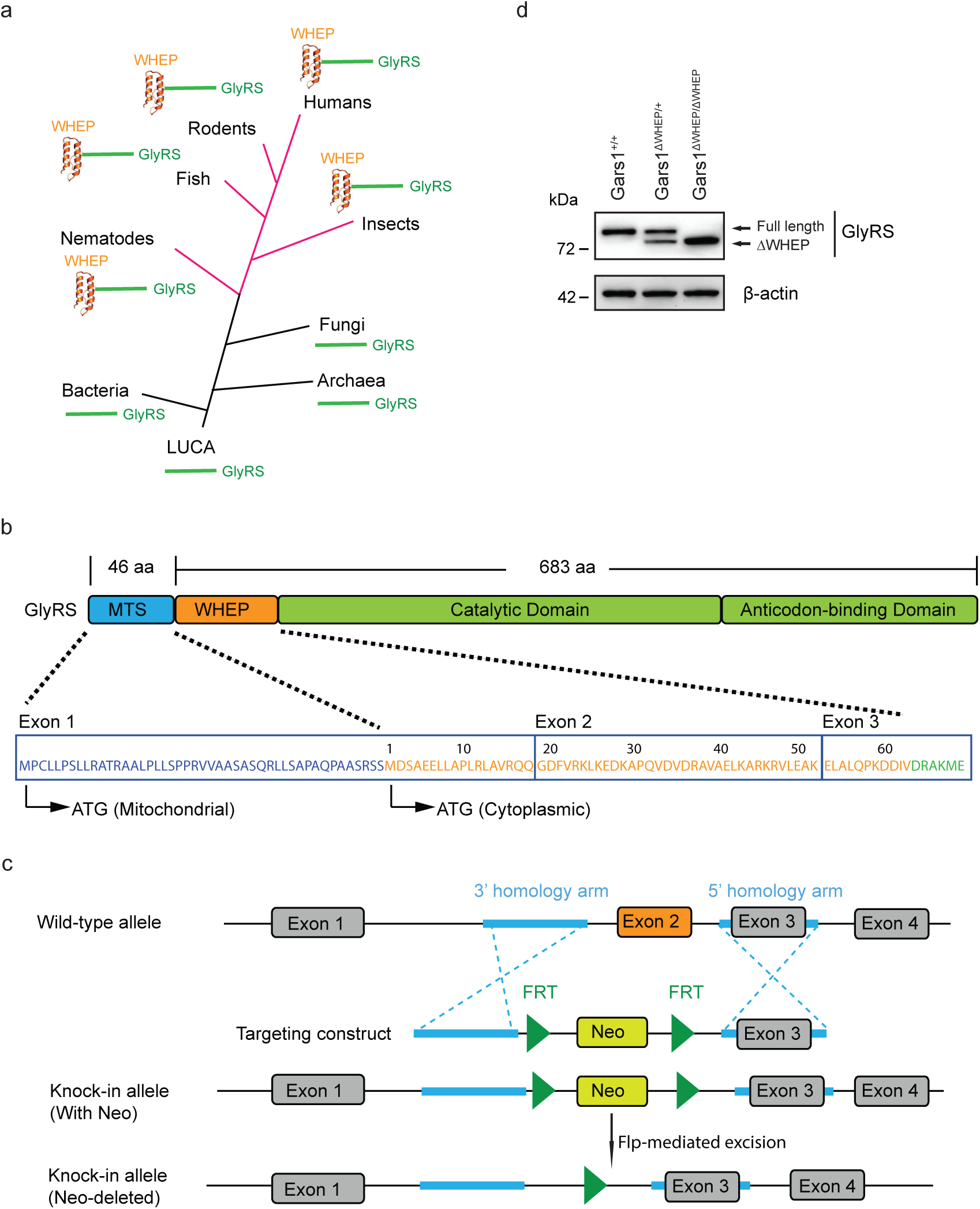
The WHEP domain of GlyRS and generation of *Gars1*^ΔWHEP^ mouse. **a.** During evolution, GlyRS WHEP domain first appeared in nematodes and has remained conserved through to humans. **b.** Mouse GlyRS domain arrangement, size, and sequence of MTS and the WHEP domain. WHEP domain is encoded across the first 3 exons of *Gars1* gene and exon 2 encodes for the majority of the domain. MTS: Mitochondrial Targeting Signal. **c.** Schematic illustration for the generation of *Gars1*^ΔWHEP^ mouse model. Exon 2 was substituted by an FRT-flanked neomycin resistance gene (NEO) cassette through homologous recombination, and the NEO cassette was subsequently removed by Flp-mediated excision after crossing with FLP-expressing C57BL/6J mice. **d.** Validation of protein expression of GlyRS^ΔWHEP^ in primary mouse embryonic fibroblasts (MEFs) isolated from WT, heterozygous or homozygous *Gars1*^ΔWHEP^ mice. The top arrow indicated full-length GlyRS protein, while the bottom arrow indicates GlyRS^ΔWHEP^ protein. A representative experiment out of five is shown.

### 2.2 *Gars1^ΔWHEP/ΔWHEP^*animals exhibit late embryonic or neonatal lethality

Inter-crossing of C57BL/6J *Gars1^ΔWHEP/+^* mice was able to produce *Gars1^ΔWHEP/ΔWHEP^* homozygous embryos close to the mendelian ratio at E12.5 with indistinguishable appearance from the littermates (Supplementary Figure S1a, b). However, at E18.5, *Gars1^ΔWHEP/ΔWHEP^* embryos had a lower (16%) than expected (25%) ratio (Supplementary Figure S1c) and significantly smaller body weights (Figure 2a). Few homozygous mutant pups were live-born (Supplementary Figure S1d), all of which struggled to breathe and died shortly after birth (Supplementary Video S1). Therefore, homozygous *Gars1^ΔWHEP/ΔWHEP^*animals exhibit late embryonic or neonatal lethality. No obvious defect was observed in the heterozygous *Gars1^ΔWHEP/+^* animals.

**Figure 2.**
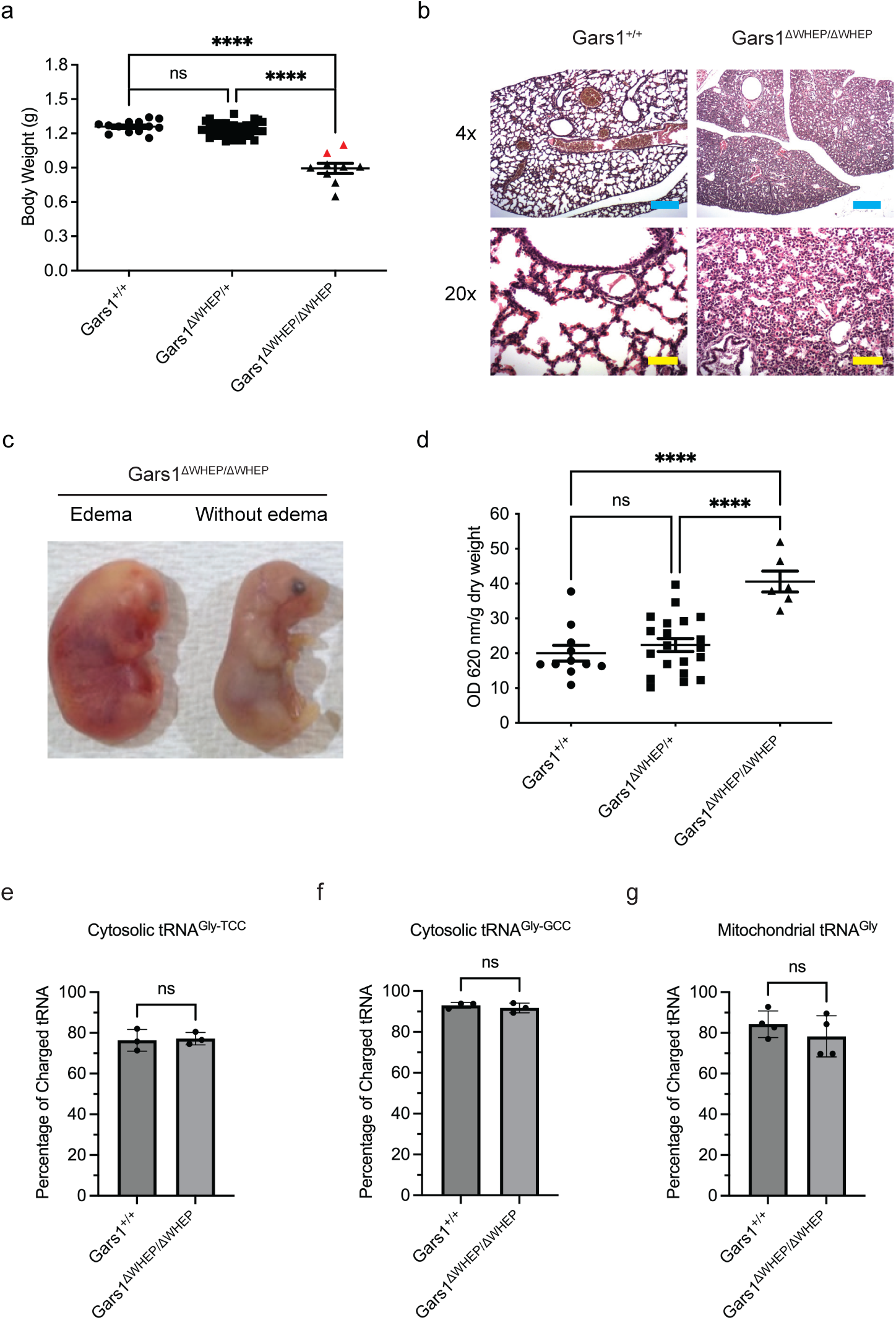
Phenotypes of homozygous *Gars1*^ΔWHEP^ mouse. **a.** Body weight of mouse embryos at E18.5. Each dot means one embryo and the red dots indicate embryos with edema. Data is represented as mean ± S.E.M of four independent experiments. Statistical analysis: ordinary one-way ANOVA with Tukey’s multiple comparisons test; ns = P > 0.05; **** P < 0.0001. **b.** Representative H&E images of lung sections from E18.5 wild-type and *Gars1*^ΔWHEP/ΔWHEP^ mouse embryos of three independent experiments. Upper panels: 4X original objective magnification, scale bar = 200 μm. Bottom panels: 20X original objective magnification, scale bar = 100 μm. **c.** Representative images of homozygous *Gars1*^ΔWHEP^ embryos at E18.5, with and without edema. **d.** Evans Blue dye extravasation assay demonstrates pulmonary vascular leakage *in vivo* using lungs from E18.5 embryos of wild-type, *Gars1*^ΔWHEP^ heterozygous and homozygous mice. Each dot means one embryo. Data is represented as mean ± S.E.M of four independent experiments. Statistical analysis: ordinary one-way ANOVA with Tukey’s multiple comparisons test; ns = P > 0.05; **** P < 0.0001. **e-g.** Quantification of acidic gel northern blot showing the *in vivo* charging status of cytosolic tRNA^Gly-TCC^ (**e**), cytosolic tRNA^Gly-GCC^ (**f**) and mitochondrial tRNA^Gly^ (**g**) from lungs of E18.5 wild-type and *Gars1*^ΔWHEP/ΔWHEP^ mouse embryos. Each dot represents one mouse. Data is represented as mean ± S.E.M of three or four independent experiments. Statistical analysis: two-tailed Student’s *t* test; ns = P > 0.05.

### 2.3 *Gars1^ΔWHEP/ΔWHEP^* lungs exhibit increased vascular permeability and are poorly inflated

In both E18.5 and P0 necropsy analyses, the lungs of *Gars1^ΔWHEP/ΔWHEP^* animals were the only organ consistently displaying a developmental delay phenotype, characterized principally by incompletely dilated airways (atelectasis) (Figure 2b). In wild-type (WT) animals, lungs exhibited dilated airways and alveoli with flattened alveolar lining cells (consistent with the expected saccular stage of pulmonary development)[<u>19</u>]. In contrast, *Gars1^ΔWHEP/ΔWHEP^* mice showed less extensive airway dilation with cuboidal alveolar lining cells (consistent with the earlier pseudoglandular stage of lung development) and scattered intraluminal cells. These morphological observations are indicative of less efficient air circulation and oxygen uptake, consistent with the breathing difficulties observed from those pups. Motor neurons in the brainstem and their processes innervating the muscles of the thoracic walls also play important roles in respiration. However, we found no significant changes of motor axon projection pattern and no obvious neuromuscular junctions (NMJs) defects in both diaphragm and limb muscles of *Gars1^ΔWHEP/ΔWHEP^*embryos (data not shown).

Interestingly, in addition to smaller sizes, about 20% of *Gars1^ΔWHEP/ΔWHEP^* embryos showed a global edema phenotype at E18.5 (Figure 2a, c). As edema can be caused by increased permeability of capillaries, we speculated that the incomplete dilation of the lung airways of *Gars1^ΔWHEP/ΔWHEP^* animals may be due to pulmonary vascular leakage. To test this hypothesis, embryos at E18.5 were isolated and injected with Evans blue dye through the temporal vein. After heart perfusion to remove residual dye within blood vessels, lungs were excised, and the amount of extravasated dye was quantified to measure vascular leakage. The *Gars1^ΔWHEP/ΔWHEP^* lungs showed a significantly higher amount of the dye compared to the control lungs, confirming pulmonary vascular leakage (Figure 2d).

### 2.4 Disruption of the WHEP domain does not affect tRNA aminoacylation in the lung

Previous *in vitro* studies suggested that deletion of the WHEP domain can slightly reduce the catalytic efficiency of GlyRS[<u>5-6</u>]. To investigate whether the lung phenotype of *Gars1^ΔWHEP/ΔWHEP^* mouse could be caused by a potential defect in tRNA aminoacylation, acidic gel northern blot was applied to examine the tRNA^Gly^ charging status in the lung. We tested both cytosolic tRNA^Gly-TCC^ and tRNA^Gly-GCC^ isoacceptors as well as mitochondrial tRNA^Gly^ and found no difference in their charging status between the *Gars1^ΔWHEP/ΔWHEP^* and the WT animals (Figure 2e, f, g, Supplementary Figure S2a, b, c). Notably, tRNA^Gly-GCC^ has a higher charging level (∼90%) than tRNA^Gly-TCC^ (∼75%), correlating with the relative abundance of the isoacceptors [<u>20</u>]. A similar result was reported from HEK293T cells [<u>21</u>]. We concluded that deletion of the WHEP domain from GlyRS does not affect tRNA charging in the lung.

### 2.5 WHEP domain is required for the release of GlyRS from cells

To understand whether the WHEP domain impacts the secretion of GlyRS, we examined the amount of soluble synthetase freely released in the culture media. GlyRS is found in the culture medium of *Gars1^+/+^* MEFs, but GlyRS^ΔWHEP^ is barely detected in the culture media of *Gars1^ΔWHEP/ΔWHEP^* and *Gars1^ΔWHEP/+^* MEFs (Figure 3a, Supplementary Figure S3a), suggesting that the WHEP domain is required for the free release of GlyRS from cells. To confirm the result in different cell types, we transfected NSC-34 motor neuron cells with plasmids encoding wild-type GlyRS or GlyRS^ΔWHEP^. Similarly, although the expression level of each construct was similar, the amount of GlyRS^ΔWHEP^ detected in cell culture medium was much less than that of the wild-type GlyRS (Figure 3b).

**Figure 3.**
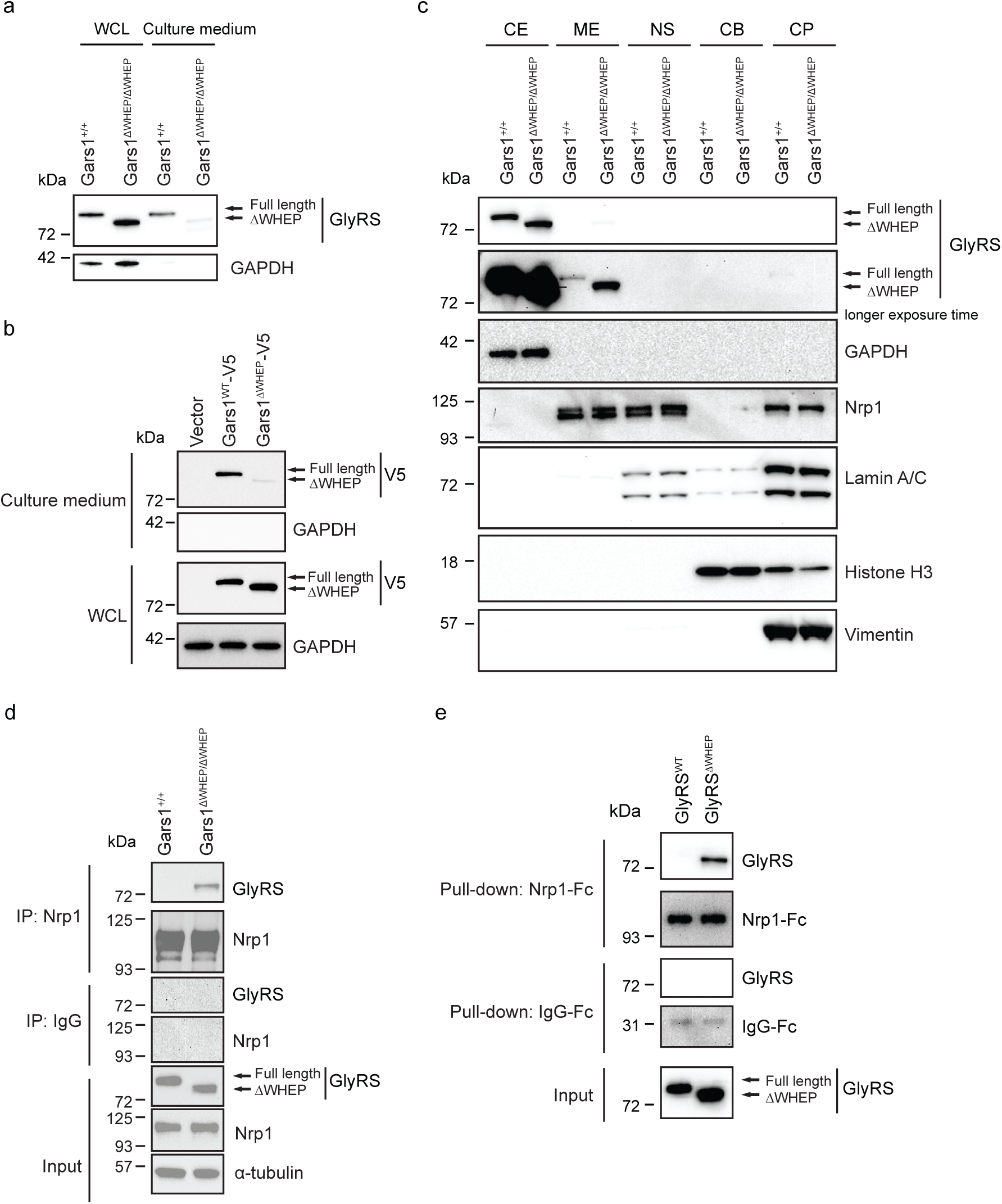
Deletion of WHEP domain affects GlyRS secretion, causes membrane retention and aberrant interaction with Nrp1. **a.** Western blot shows GlyRS protein level from whole cell lysate (WCL) and culture medium of wild-type and *Gars1*^ΔWHEP/ΔWHEP^ MEFs. A representative experiment out of three is shown. **b.** Western blot shows GlyRS protein level from WCL and culture medium of NSC-34 cells transfected with plasmids encoding Gars1^WT^-V5, Gars1^ΔWHEP^-V5, and vector control. A representative experiment out of three is shown. **c.** Subcellular protein fractionation of wild-type and *Gars1*^ΔWHEP/ΔWHEP^ MEFs. CE: cytosol extraction, ME: membrane extraction, NS: nuclear soluble, CB: chromatin-bound, CP: cytoskeletal proteins. A representative experiment out of three is shown. **d.** Co-immunoprecipitation (Co-IP) of endogenous Nrp1 and GlyRS^ΔWHEP^ from lysates of cultured MEFs. A representative experiment out of three is shown. **e.** *In vitro* pull-down experiments further confirm GlyRS^ΔWHEP^ as a Nrp1 binding partner. Recombinant Nrp1 (C-terminus Fc tagged) protein was bound to Dynabeads Protein G and then incubated with purified recombinant GlyRS^WT^ or GlyRS^ΔWHEP^ proteins. A representative experiment out of three is shown.

### 2.6 GlyRS^ΔWHEP^ was enriched in membrane fraction where Nrp1 is also detected

Disrupting the WHEP domain diminishes the release of GlyRS from cells but not its expression. Therefore, we investigated the subcellular localizations of GlyRS^ΔWHEP^. Contents in subcellular compartments were sequentially extracted in the order of cytosol (CE), membrane (ME), and nucleus. The nucleus fraction was further separated into nuclear soluble (NS), chromatin-bound (CB), and cytoskeletal proteins (CP) fractions. Glyceraldehyde-3-phosphate dehydrogenase (GAPDH), Nrp1, Lamin A/C, Histone H3 and Vimentin were used as markers for CE, ME, NS, CB, and CP fractions, respectively (Figure 3c). GlyRS proteins were detected mainly in the cytoplasmic fraction, with a small percentage localized in the membrane fraction. Interestingly, GlyRS^ΔWHEP^ protein showed an increased abundance in the membrane fraction (Figure 3c, Supplementary Figure S3b), which is the only subcellular compartment where both GlyRS and Nrp1 are detected, prompting us to study the impact of the WHEP domain on GlyRS-Nrp1 interaction.

### 2.7 Disrupting WHEP domain enhances GlyRS-Nrp1 interactions *in cellulo* and *in vitro*

We studied the impact of the WHEP domain on GlyRS-Nrp1 interaction both *in vitro* with a pull-down experiment and *in cellulo* with co-immunoprecipitation. MEFs abundantly express Nrp1 and GlyRS (Figure 3d). The GlyRS-Nrp1 interaction was clearly detected in *Gars1^ΔWHEP/^*^ΔWHEP^ but not in *Gars1^+/+^* MEFs (Figure 3d). Consistently, pull-down of recombinant GlyRS proteins with the C-terminally Fc-tagged extracellular region of Nrp1 showed that GlyRS^ΔWHEP^ has a much stronger interaction with Nrp1 compared to its WT counterpart (Figure 3e), suggesting that disrupting the WHEP domain enhances the extracellular GlyRS-Nrp1 interaction and that the GlyRS^ΔWHEP^-Nrp1interaction is direct.

### 2.8 Disrupting the WHEP domain in *Gars1* stimulates Nrp1 internalization

We also probed the functional impact of disrupting the GlyRS WHEP domain on Nrp1. One of the key functions of Nrp1 is mediating endocytosis, a process that can be regulated by various ligands, including oligo-guanosine nucleotides [<u>22</u>]. Indeed, an 18-mer of deoxyguanosine (G18) induced Nrp1 internalization in MEFs, as measured by the decrease of cell surface Nrp1 (Figure 4a). Interestingly, compared to the WT cells, the G18-stimulated Nrp1 internalization was enhanced in *Gars1^ΔWHEP/^*^ΔWHEP^ MEFs (Figure 4a), suggesting that the aberrant GlyRS^ΔWHEP^-Nrp1 interaction further promotes G18-induced Nrp1 internalization.

**Figure 4.**
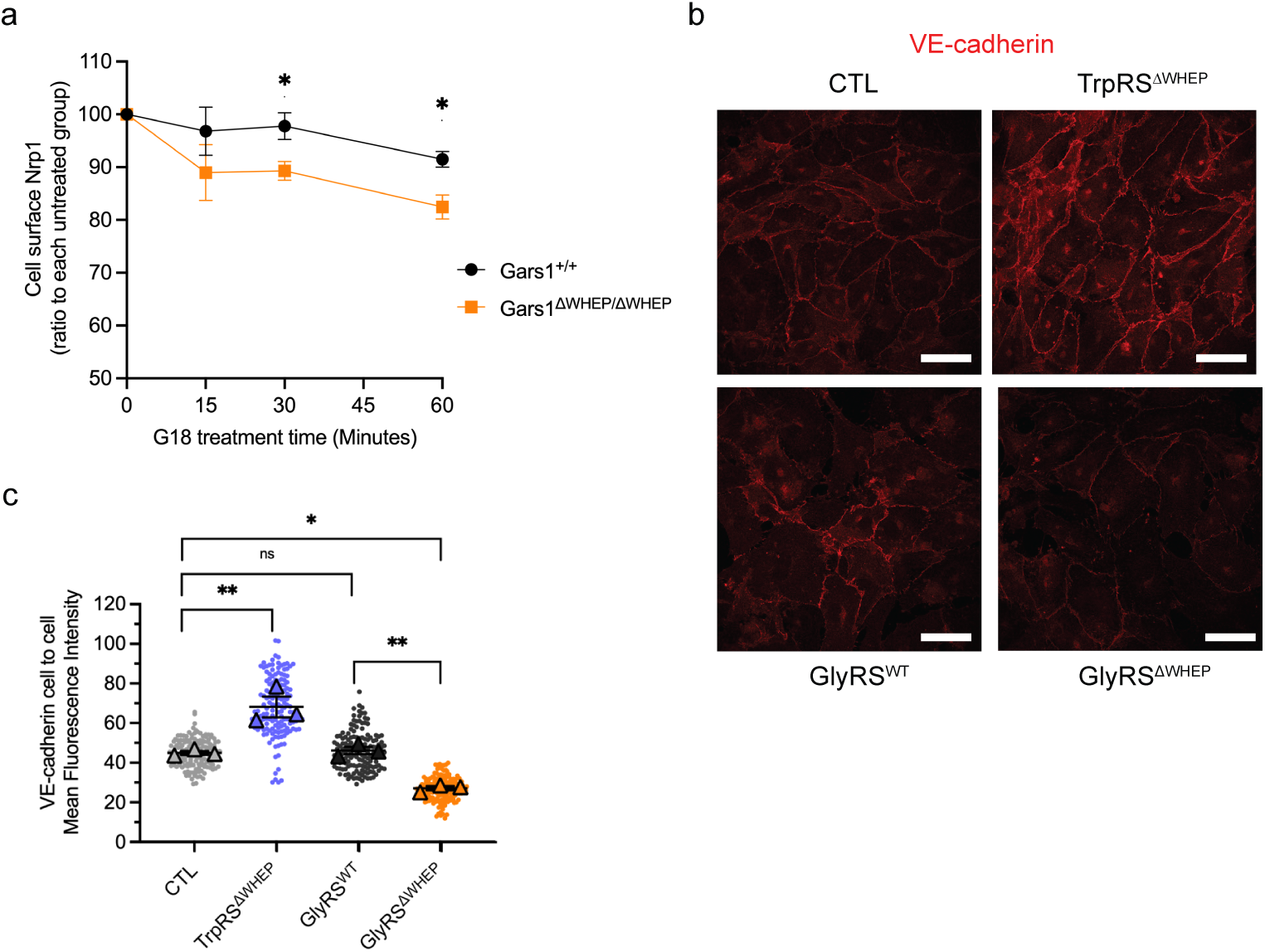
GlyRS^ΔWHEP^ facilitates Nrp1 internalization and reduces VE-cadherin accumulation at the intercellular contacts. **a.** Cell surface Nrp1 is detected by FACS after G18 stimulation for different time points in wild-type and *Gars1*^ΔWHEP/ΔWHEP^ MEFs. Data is represented as mean ± S.E.M of three independent experiments. Statistical analysis: two-way ANOVA with the Geisser-Greenhouse correction; ns = P = 0.41; * P = 0.045, * P = 0.028. **b.** Cell-cell contact VE-cadherin is visualized by confocal microscopy using HUAECs transduced with GlyRS^WT^, GlyRS^ΔWHEP^, TrpRS^ΔWHEP^ or control lentivirus (CTL). 63X original objective magnification, scale bar = 10 μm. **c.** Quantification of Figure 4b. Data are the mean +/-SEM of three biologically independent experiments (depicted in different color shades, VE-Cadherin at AJs evaluated in 50 cells in each sample in each of the three independent biological experiments). For statistical evaluation, parametric two-tailed ANOVA with Bonferroni correction was applied; ns = P > 0.05; * P ≤ 0.05; ** P ≤ 0.01.

### 2.9 GlyRS^ΔWHEP^ stimulates VE-cadherin turnover in endothelial cells

Our previous work showed that deletion of the WHEP domain from TrpRS also enhances the Nrp1 interaction [<u>8</u>]. However, TrpRS^ΔWHEP^ (mini-WARS) attenuates the endocytic function of Nrp1. When overexpressed in human umbilical artery endothelial cells (HUAECs), TrpRS^ΔWHEP^ acts as an inhibitor of Nrp1-mediated VE-cadherin turnover, leading to increased VE-cadherin accumulation at sites of the intercellular contact and decreased vascular permeability [<u>8</u>]. To clarify whether the function of GlyRS^ΔWHEP^ resembles or opposes that of TrpRS^ΔWHEP^, we directly compared the impact of GlyRS^ΔWHEP^ and TrpRS^ΔWHEP^ on VE-cadherin localization at endothelial adherens junctions. Compared with HUAECs transduced with control lentivirus (CTL) or full-length GlyRS, HUAECs overexpressing GlyRS^ΔWHEP^ showed attenuation of VE-cadherin-positive intercellular contacts, an opposite effect of TrpRS^ΔWHEP^ (Figure 4b and 4c, Supplementary Figure S4). Therefore, although deletion of the WHEP domain activates the Nrp1 interaction for both TrpRS and GlyRS, the functional impact is the opposite. Expression of GlyRS^ΔWHEP^ stimulates Nrp1-mediated VE-cadherin removal from endothelial intercellular contact, which in turn would enhance vascular permeability, thus corroborating our *in vivo* observations of pulmonary edema in *Gars1^ΔWHEP/ΔWHEP^* mice.

### 2.10 Genetic knockdown of Nrp1 partially rescues *Gars1^ΔWHEP/^*^ΔWHEP^ phenotypes

To further evaluate if Nrp1 is relevant to the *Gars1^ΔWHEP/^*^ΔWHEP^ phenotypes in *vivo,* we tested the genetic interaction between *Gars1*^ΔWHEP^ and Nrp1. Because the aberrant GlyRS^ΔWHEP^-Nrp1 interaction stimulates Nrp1, at least for its endocytic activity, we speculated that knockdown of Nrp1 may have a positive impact on *Gars1^ΔWHEP/^*^ΔWHEP^ animals. Nrp1 knockout (Nrp1^-/-^) embryos are not viable [<u>13</u>], so we intercrossed *Gars1^ΔWHEP/^*^+^ and Nrp1^+/-^ animals. Knockdown of Nrp1 showed decreased expression of Nrp1 in both spinal cord and lung from *Gars1^ΔWHEP/^*^+^; Nrp1^+/-^ mice (Supplementary Figure S5a). The knockdown did not impact the viability of *Gars1^ΔWHEP/^*^ΔWHEP^ embryos, and *Gars1^ΔWHEP/^*^ΔWHEP^; Nrp1^+/-^ animals still die late in gestation or shortly after birth. Nevertheless, the double mutant embryos show increased body weights (Figure 5a) and decreased pulmonary vascular leakage compared to *Gars1^ΔWHEP/^*^ΔWHEP^ mice (Figure 5b), suggesting that the aberrant GlyRS^ΔWHEP^-Nrp1 interaction is partially responsible for the phenotypes of the *Gars1^ΔWHEP/ΔWHEP^* animals. It is worth noting that the Nrp1 expression level is remarkably high in the lung (Supplementary Figure S5b). Possibly, the level of Nrp1 knockdown is insufficient to offset the aberrant activation by *Gars1^ΔWHEP^*.

**Figure 5.**
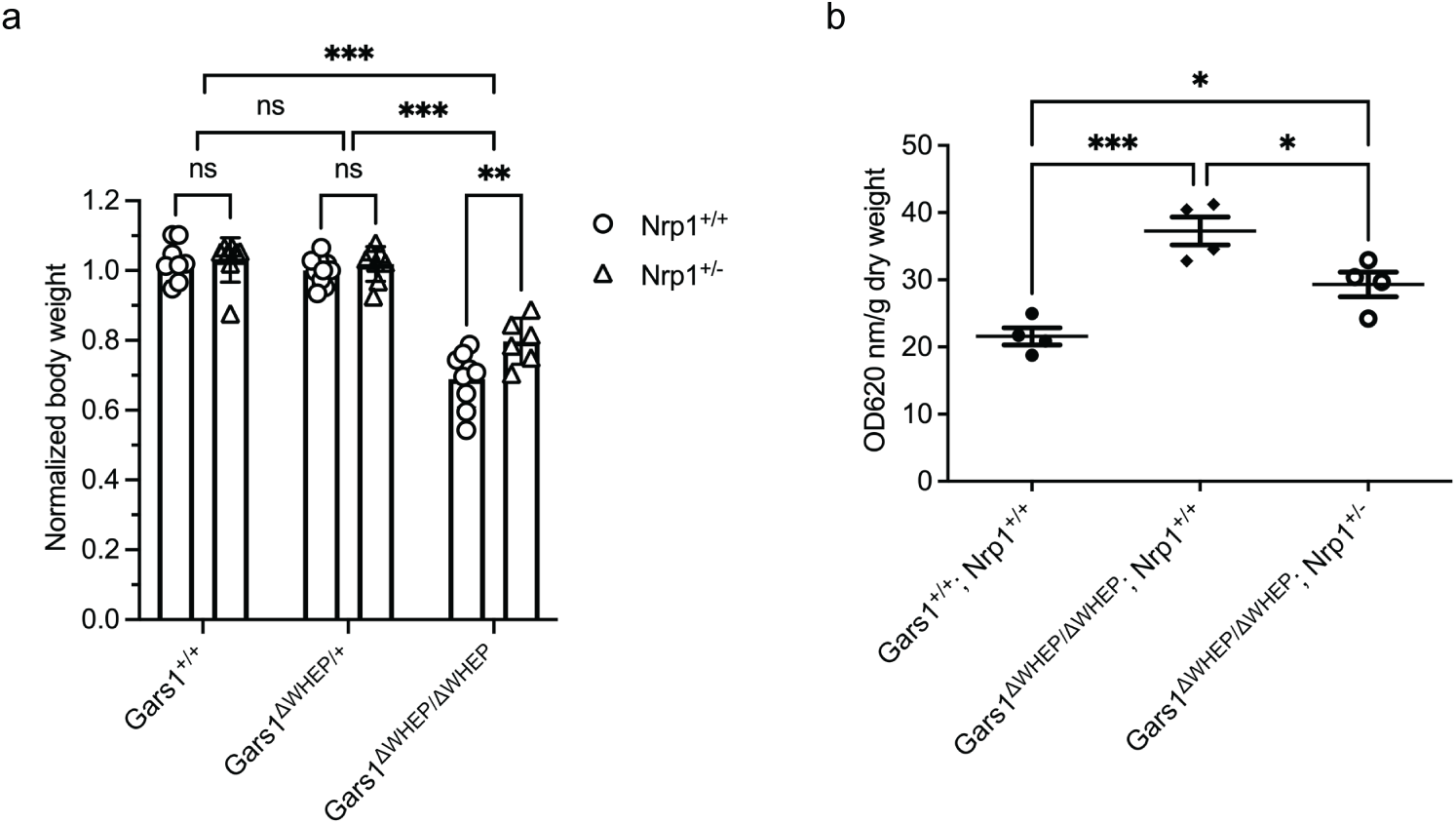
Genetic knockdown of Nrp1 partially rescues *Gars1*^ΔWHEP/ΔWHEP^ mouse phenotypes. **a.** Normalized body weight of E18.5 embryos from crosses between *Gars1*^ΔWHEP/+^ and *Gars1*^ΔWHEP/+^; Nrp1^+/-^ mice. Data is represented as mean ± S.E.M of five independent experiments. Statistical analysis: two-way ANOVA with the Šidák correction; ns = P > 0.05; **p≤0.01, ***p≤0.001. **b.** Evans Blue dye extravasation assay comparing pulmonary vascular leakage using lungs from E18.5 embryos of *Gars1*^ΔWHEP/ΔWHEP^; Nrp1^+/+^, *Gars1*^ΔWHEP/ΔWHEP^; Nrp1^+/-^, and wild-type mice. Data is represented as mean ± S.E.M of three independent experiments. Statistical analysis: two-way ANOVA with the Tukey’s correction; *p≤0.05, ***p≤0.001.

## 3. Discussion

*Gars1^ΔWHEP/ΔWHEP^* animals exhibit late embryonic or neonatal lethality (Supplementary Figure S1c, 1d). In the rare instances of live-born homozygous mutant pups, they rapidly succumbed after birth, displaying gasping (Supplementary Video S1). Consistently, our necropsy analyses revealed that the most consistent phenotypes occur in the lung, where *Gars1^ΔWHEP/ΔWHEP^*animals exhibited an obvious developmental delay characterized by airways lined by cuboidal epithelium and reduced dilation of airways at all levels (Figure 2b). Moreover, lung parenchyma (alveolar walls) is richly vascularized, and the Evans Blue dye extravasation assay demonstrated significant extravascular dispersion of dye (i.e., vascular leakage) in E18.5 *Gars1^ΔWHEP/ΔWHEP^* lungs (Figure 2d). Some embryos also presented with whole-body edema (Figure 2c). Taken together, the vascular leakage in the lung and whole-body edema suggest that the mutation may interfere with endothelial barrier function or cell-cell junctions, perhaps to a differing degree depending on the organ and vascular bed.

In our previous work, GlyRS, along with two other WHEP-domain containing aaRSs (i.e., TrpRS and HisRS), was found to interact with Nrp1 in endothelial cells [<u>8</u>]. Nrp1 plays a significant role in the regulation of vascular permeability, in part through its endocytosis function and its interaction with adhesion receptors like VE-cadherin and platelet and endothelial cell adhesion molecule 1 (PECAM1), which facilitate the internalization of VE-cadherin from the cell surface and destabilize the endothelial cell-cell junctions [<u>8</u>]. Interestingly, here we find the GlyRS^ΔWHEP^ protein to be trapped in the membrane fraction of MEFs, co-localizing with Nrp1 (Figure 3c). We also find, both in cells and *in vitro*, that GlyRS^ΔWHEP^ makes a stronger interaction with Nrp1 compared with the full-length GlyRS (Figure 3d, 3e). Compared to WT cells, G18-stimulated Nrp1 internalization was enhanced in *Gars1^ΔWHEP/^*^ΔWHEP^ MEFs (Figure 4a), suggesting the aberrant GlyRS^ΔWHEP^-Nrp1 interaction may over-activate the endocytic function of Nrp1 and induce vascular leakage. Indeed, the vascular leakage is partially rescued in *Gars1* ^ΔWHEP/ΔWHEP^ mice when Nrp1 is heterozygously deleted (Figure 5b).

How can the GlyRS^ΔWHEP^ -Nrp1 interaction stimulate the endocytic function of Nrp1? This question is especially interesting considering that GlyRS^P234KY^, a CMT2D-causing mutant form of GlyRS found in our previous work, also gains a stronger interaction with Nrp1 than the WT GlyRS. Interestingly, GlyRS^P234KY^ antagonizes the VEGF-Nrp1 interaction without affecting the vasculature [<u>7b</u>, <u>23</u>]. Rather, GlyRS^P234KY^ inhibits the neural protective role of VEGF-Nrp1 signaling while the motor function deficit of the *Gars1*^P234KY/+^ CMT2D mouse model is exacerbated with a heterozygous Nrp1 knockout [<u>7b</u>].

Therefore, GlyRS^P234KY^ differentiates from GlyRS^ΔWHEP^ for their impact on Nrp1 function and the systems involved. An opposite impact was also observed between TrpRS and GlyRS (Figure 4b, 4c), with both gaining Nrp1 interaction when their WHEP domain is removed but with TrpRS^ΔWHEP^ acting as a negative rather than a positive regulator for Nrp1-mediated VE-cadherin turnover, thereby causing a decrease in vascular permeability [<u>8</u>].

Consistent with what has been shown in endothelial cells [<u>22</u>], oligo-guanosine G18 can also weakly stimulate Nrp1 internalization in wild-type MEFs (Figure 4a). This effect is enhanced in *Gars1^ΔWHEP/^*^ΔWHEP^ mutant cells (Figure 4a). Considering MEFs express a high level of Nrp1 but a low level of VEGFR2 (Supplementary Figure S6), the activation effect of *Gars1^ΔWHEP^* on Nrp1 internalization is likely to be co-receptor independent. Indeed, Roth et al. have shown that the Nrp1-mediated vascular permeability can occur independently of VEGFR2 activation and that ligand-induced Nrp1 clustering or multimerization at cell-cell contacts is necessary for Nrp1-mediated endocytosis and vascular permeability induction[<u>16a</u>]. Most aaRSs, including GlyRS and TrpRS, function as a dimer during tRNA aminoacylation. The intact tRNA charging activity in *Gars1^ΔWHEP/ΔWHEP^* tissues suggests that disruption of the WHEP domain does not negatively impact dimerization. In contrast, many mutations causing Charcot-Marie-Tooth disease are located near the dimer interface of GlyRS [<u>24</u>], and the mutations alter the dimer interface [<u>10</u>]. For example, GlyRS^P234KY^ has been shown to have a weakened dimer association [<u>7b</u>]. Nrp1 itself forms dimers through its transmembrane and juxtamembrane c domains. Bridging the Nrp1 dimers through dimers of aaRS could cluster the receptor. We speculate that both the capacity to form dimers and the specific configuration of the dimers could impact the ability of an aaRS to multimerize Nrp1. TrpRS^ΔWHEP^ likely also form dimers, but the configuration may not be compatible with clustering of Nrp1. Instead, TrpRS^ΔWHEP^, like GlyRS^P234KY^, may competitively bind to Nrp1 and prevent the receptor from being activated by other ligands. Further research is needed to clarify these mechanisms.

The *Gars1^+/^*^ΔWHEP^ animals have no obvious phenotypes even though the GlyRS^ΔWHEP^ protein is also trapped in the membrane fractions (Supplementary Figure S3a, 3b) and presumably can interact with Nrp1. There are two perspectives to consider in the explanation. On one hand, the level of GlyRS^ΔWHEP^ in the membrane may not have reached the threshold of toxicity; on the other hand, the remaining of GlyRS^WT^ in the circulation of *Gars1^+/^*^ΔWHEP^ animals may provide some protective effect that is completely missing in the *Gars1^ΔWHEP/^*^ΔWHEP^ animals. Extracellular functions have been reported for GlyRS. For example, extracellular vesicle-associated GlyRS, through its WHEP domain, interacts with cadherin EGF LAG seven-pass G-type receptor 2 (CELSR2) to activate M1 macrophage polarization and phagocytosis as well as to suppress tumor initiation[<u>25</u>]. The potential protective function of extracellular GlyRS relevant to endothelial cells remains to be clarified.

It is worth noting that a natural splice variant of HisRS was found to constitute only the WHEP domain tailed with a C-end-rule peptide (K^57^TPK^60^) [<u>26</u>]. The HisRS WHEP domain can bind to Nrp2, but not Nrp1, and exhibits immune-suppressive activity [<u>7c</u>, <u>27</u>]. As mentioned above, TrpRS^ΔWHEP^ can also occur naturally as a splice variant. These findings highlight the aaRS family as a novel class of Nrp1 regulators. Similar splice variants have not yet been identified for GlyRS, but considering that the WHEP domain of GlyRS is appended to the corresponding enzymatic core through a flexible linker [<u>4</u>], it is conceivable that GlyRS^ΔWHEP^ may occur naturally, for example, through proteolysis. Therefore, our *Gars1^ΔWHEP/^*^ΔWHEP^ animals may represent an over-activated natural regulation, revealing a physiological role of GlyRS in regulating blood vessel biology.

## 4. Experimental Section

### DNA Constructs

Mouse GlyRS^WT^ and mouse GlyRS^ΔWHEP^ constructs were cloned into pcDNA6/V5-His C vector (ThermoFisher Scientific, catalog # V22020). Mouse GlyRS^WT^, mouse GlyRS^ΔWHEP^ and human TrpRS^ΔWHEP^ were subcloned into a third-generation lentiviral vector pCCL.sin.cPPT.PGK.GFP.WPRE[<u>28</u>] (pCCL), with In-Fusion 2.0-CF Dry-DownPCR Cloning-Kit (catalog no. 639607; Clontech Laboratories, Mountain View, CA).

### Animal Models

The *Gars1*^ΔWHEP^ mice were generated by deletion of exon 2 of the *Gars1* gene. An FRT-flanked Neo cassette was introduced in exon 2 that was later bred out to create the Deletion exon 2-Neo strain. The Neo was removed through Flp-mediated excision to create the *Gars1*^ΔWHEP^ strain. The F0 mice were crossed with wild-type C57BL/6J mice to get *Gars1*^ΔWHEP/+^ mice that were used for breeding. Mouse experiments were approved in advance by and conducted in accordance with the guidelines of the Scripps Research IACUC. For the genetic interaction study, *Gars1*^ΔWHEP/+^ mice were crossed with Nrp1^+/-^ mice.

### Cell Culture

Mouse motor neuron cell line NSC-34 was purchased from ATCC. This cell line was grown in DMEM (Dulbecco’s Modified Eagle Medium, Gibco) with 10% heat-inactivated FBS (fetal bovine serum, Gibco) and 1% penicillin/streptomycin (Gibco). Cells were cultured at 37°C in a humidified incubator with 5% CO_2_.

Primary MEFs were isolated from embryos from pregnant (*Gars1*^ΔWHEP/+^ x *Gars1*^ΔWHEP/+^) females. All dissecting tools were washed in 70% ethanol and then put under UV lamp in a TC hood. Embryos were collected between E12.5 – E14.5 from timed pregnant females, washed in 70% ethanol, and then placed in a petri dish with PBS with 1% penicillin/streptomycin. The embryos were separated from placenta; the head, inner organs, and limbs were removed; and the remaining blood was rinsed off in PBS with 1% penicillin/streptomycin. The remaining parts of embryos were minced with razor blades in a 100 mm petri dish, and then 2 mL 0.25% Trypsin-EDTA was added before incubating at 37°C in a cell incubator for 10 minutes with intermittent shaking. After 10 minutes, 4 mL growth media containing DMEM with 10% embryonic stem cell FBS (ThermoFisher Scientific, 16141079) and 1% penicillin/streptomycin was added. Cells in growth media were transferred to a 15 mL conical tube and allowed to settle. The cell suspension was aspirated and transferred to a new 15 mL conical tube. Cells were spun at 200 x *g* for 2 minutes, supernatant was removed. The cell pellet was resuspended in 10 mL growth media, and then cells were plated in a 10 cm dish. This preparation represents passage 0. Primary MEFs were cultured at 37°C in a humidified incubator with 5% CO_2_. All experiments were conducted at passages 1-3.

Primary human ECs were previously described[<u>8</u>]. Briefly, umbilical arteries were cannulated with a blunt 17-gauge needle that was secured by clamping the cord over the needle. The umbilical arteries were then perfused with 50 ml of phosphate-buffered saline (PBS) to wash out the blood. Next, 10 ml of 0.2% collagenase A (Roche Diagnostics, # 11088793001,) diluted in cell culture medium were infused into the umbilical arteries and incubated 30 min at room temperature. The collagenase solution containing the ECs was flushed from the cord by perfusion with 40 ml of PBS, collected in a sterile 50 ml centrifuge tube, and centrifuged 5 min at 800 x g. Cells were first resuspended in Endothelial Cell Growth Basal Medium (EBM-2) supplemented with EGM-2 BulletKit (Lonza) (EGM-2), and subsequently plated in cell culture dishes that had been previously adsorbed with 1% gelatin from porcine skin (G9136, Sigma-Aldrich).

### Antibodies and Reagents

Mouse monoclonal antibody (mAb) anti-GlyRS (Homemade, 12G1, for western blot-WB 1:3000); Mouse mAb anti-ϕ3-actin (Cell Signaling, 8H10D10/#3700, WB 1:1000); Rabbit mAb anti-GAPDH (Cell Signaling, 14C10/#2118, WB 1:1000); Mouse mAb anti-V5 Tag (ThermoFisher Scientific, R960-25, WB 1:2000); Rabbit mAb anti-Nrp1 (Abcam, ab81321, WB 1:2000); Mouse mAb anti-Lamin A/C (Cell Signaling, 4C11/#4777, WB 1:2000); Rabbit mAb anti-Histone H3 (Cell Signaling, #9715, WB 1:1000); Rabbit mAb anti-Vimentin (Cell Signaling, D21H3/#5741, WB 1:1000); Rabbit polyclonal anti-Na, K-ATPase (Cell Signaling, #3010, WB 1:1000); rabbit polyclonal anti-TrpRS (ThermoFisher Scientific, PA5-29102, WB 1:1000); Mouse mAb anti-vinculin (Sigma-Aldrich, V9131, WB 1:2000); Goat polyclonal anti-VE-cadherin (Santa Cruz Biotechnology, C-19/sc-6458, IF 1:200); Rabbit polyclonal anti-α-tubulin (Cell Signaling, #2144, WB 1:1000); Goat anti-rabbit IgG, HRP-linked secondary Ab (Cell Signaling, #7074, WB 1:5000); Horse anti-mouse IgG, HRP-linked secondary Ab (Cell Signaling, #7076, WB 1:5000); Alexa Fluor-555 donkey anti-goat IgG (H+L) secondary Ab (Life Technologies, A21432, IF 1:400); Rat IgG_2A_ anti-mouse Nrp-1 Alexa Fluor® 647-conjugated mAb (R&D, FAB59941R, FACS 1:100); Rat IgG_2A_ Alexa Fluor® 647-conjugated isotype control (R&D, IC006R, FACS 1:100); Rat IgG_2a_ kappa anti-VEGFR2 (CD309/FLK1) mAb, PE (ThermoFisher Scientific, #12-5821-82, FACS 1:100); Rat IgG_2a_ kappa isotype control, PE (ThermoFisher Scientific, #12-4321-80, FACS 1:100); Goat anti-human IgG Fc HRP preadsorbed (abcam, ab98624, WB 1:10000); Dynabeads protein G beads (ThermoFisher Scientific, 10003D).

### Recombinant Proteins

Recombinant human Nrp1-Fc chimera protein (R&D Systems, 10455-N1); the control human IgG Fc protein (R&D Systems, 110-HG). For the expression and purification of recombinant mouse GlyRS isoforms, DNA encoding mouse GlyRS^WT^ or GlyRS^ΔWHEP^ with a N-terminal His6-SUMO tag was cloned into pET-28a (+) vector (Novagen). GlyRS expression was induced in *E. coli* BL21 (DE3) cells with 1 mM Isopropyl β-D-1-thiogalactopyranoside (IPTG). Following cell lysis, GlyRS was purified using Ni-NTA beads (Qiagen). The N-terminal His6-SUMO tag was cleaved with sumo protease ULP1. The resulting tag-free GlyRS proteins were further purified using HiLoad 16/60 Superdex 200 prep grade column (GE Healthcare).

### Cell Transfection

When NSC-34 cells had reached 70-80% confluence, transfections were conducted using Lipofectamine 3000 (Life Technology) according to the manufacturer’s instructions. NSC-34 cells were transfected with mouse *Gars1*^WT^ or mouse *Gars1*^ΔWHEP^ constructs in pcDNA6/V5-His C vector, with the vector as control.

### Protein Secretion Assay

When transfected NSC-34 cells or primary MEFs had reached 70-80% confluence, cells were washed with Dulbecco’s PBS, no calcium, no magnesium (DPBS, ThermoFisher Scientific, #14190144) and further cultured in serum-free DMEM overnight. The culture medium was collected and centrifuged at 500g for 10 minutes at 4°C. The supernatant was transferred to a new tube and further centrifuged at 20,000g for 15 minutes at 4°C to remove contaminants. Proteins were precipitated from the supernatants with a final concentration of 10% trichloroacetic acid (TCA, Sigma #T6399, make 100% stock with dH_2_O) for 1 hour at 4°C and then centrifuged at 20,000g for 15 min at 4°C. The supernatant was carefully removed, and the pellets were washed with ice-cold acetone. The supernatant was removed after centrifuge at 20,000g for 15 min at 4°C, and the pellet was air-dried for 3 minutes. The pellet was dissolved in 2xSDS sample buffer, boiled for 10 min in 95°C and ready for western blot analysis.

### Subcellular Fractionation

Subcellular Protein Fractionation Kit for Cultured Cells (Thermo Scientific #78840) was used to segregate proteins from five different cellular compartments. Primary MEFs at 70-80% confluence were used for subcellular fractionation in accordance with the manufacturer’s instructions. Briefly, the first kit reagent, when added to a cell pellet, causes selective membrane permeabilization, releasing soluble cytoplasmic contents. The second reagent dissolves plasma, mitochondria and ER-golgi membranes but does not solubilize the nuclear membranes. After recovering intact nuclei by centrifugation, a third reagent yields the soluble nuclear extract. An additional nuclear extraction with micrococcal nuclease is performed to release chromatin-bound nuclear proteins. The recovered insoluble pellet is then extracted with the final reagent to isolate cytoskeletal proteins.

### Western Blot Assay

Cells at 70-80% confluence were washed in cold DPBS, no calcium, no magnesium (ThermoFisher Scientific, #14190144) and lysed in cold 1x RIPA buffer (ThermoFisher Scientific, #89901) with Halt^TM^ protease inhibitor cocktail (ThermoFisher Scientific, #78429) for 15 minutes on ice, and centrifuged for 10 minutes at 18,000 x *g* at 4 °C. The supernatant was collected, and the same volume of 2xSDS sample buffer was added. After boiling for 10 minutes in 95°C, the samples were loaded in protein gels and separated by SDS-PAGE, after which the proteins were transferred to PVDF membrane for immunoblotting with antibodies for protein of interest.

### In Vitro Pull-down Assay

Equal amount (about 0.5 μg) of recombinant Fc-tagged whole extracellular portion of human Nrp1 protein (Nrp1-Fc, R&D Systems, 10455-N1) or the control human IgG Fc protein (R&D Systems, 110-HG) were bound to 5 uL Dynabeads Protein G (ThermoFisher Scientific, 10003D) for 30 min at 4 °C. Equal amount (about 2 μg) of purified recombinant mouse GlyRS^WT^ or GlyRS^ΔWHEP^ protein was added to the Dynabeads coated with Nrp1-Fc or Fc control and incubated for 4 hours at 4°C. The beads were washed three times with the wash buffer (25 mM Tris-HCl, pH 7.4, containing 150 mM NaCl, 1 mM EDTA, 1% NP-40 and 5% glycerol) to remove unbound proteins. 50 μL 2xSDS sample buffer was then added to the beads to elute bound proteins and then boiled for 10 minutes in 95°C. The amount of Nrp1-Fc and GlyRS was analyzed by western blot analysis using goat anti-human IgG Fc (HRP) antibody (abcam, ab98624, 1:10000) and home-made mouse anti-GlyRS mAb (12G1, 1:3000), respectively.

### Co-Immunoprecipitation (Co-IP)

MEFs (*Gars1*^+/+^ and *Gars1*^ΔWHEP/ΔWHEP^) were seeded in 150 mm dishes. When around 80% confluent, MEFs were cross-linked with the final concentration of 0.1% PFA by adding 250 μL 4% formaldehyde (Paraformaldehyde/PFA, ThermoFisher Scientific, J61899-AK) per 10 mL media, and then incubated for 10 min at 37°C. The reaction was quenched by adding glycine to the final concentration of 125 mM. Cells were washed with cold DPBS (ThermoFisher Scientific, #14190144) and then lysed in 1.5 mL cold Pierce™ IP Lysis Buffer (ThermoFisher Scientific, 87787) with Halt^TM^ protease inhibitor cocktail (ThermoFisher Scientific, #78429). Cellular lysates were incubated for 15 min on ice, and then centrifuged for 10 minutes at 18,000 x *g* at 4 °C. The supernatant was transferred to a new Eppendorf tube (prechilled). The total protein amount was determined using the bicinchoninic acid (BCA) assay (Pierce). Equal amount (about 1 μg) of rabbit anti-Nrp1 mAb (abcam, ab81321) and the normal rabbit IgG control (abcam, ab172730) were bound to 5 μL Dynabeads Protein G (ThermoFisher Scientific) in 1mL IP buffer by rotating at room temperature for 1 hour. After 1 hour incubation, the antibody-dynabeads conjugates were separated by a magnetic stand to aspirate the supernatant before resuspending the beads in 100 μL new IP buffer. Equivalent amounts (About 1-2 mg) of total protein from cell lysates of MEFs were added to the Dynabeads coated with anti-Nrp1 or the IgG control and incubated overnight at 4°C. The beads were washed three times with IP buffer to remove unbound proteins. 50 μL 2xSDS sample buffer was then added to the beads to elute bound proteins and the sample was boiled for 20 minutes in 95°C. The amount of Nrp1 and GlyRS was analyzed by western blot analysis using rabbit anti-Nrp1mAb (abcam, ab81321) and home-made mouse anti-GlyRS mAb (12G1), respectively.

### Hematoxylin and Eosin (H&E) Staining

E18.5 embryos or newborn pups at P0 from *Gars1*^ΔWHEP/+^ intercross were harvested, dispatched by exsanguination, fixed by immersion in 4% methanol-free formaldehyde (Paraformaldehyde/PFA, ThermoFisher Scientific, J61899-AK) overnight in a cold room. After cold PBS wash, the animals were dehydrated by sequential incubations in 30%, 50%, 70%, 95% and 100% ethanol. After clearing with xylene, the animals were infiltrated with and embedded in paraffin wax. Whole embryo sagittal sections at 5 µm were stained routinely with hematoxylin and eosin. All digital images were examined and captured using Leica AT2 whole slide scanner (brightfield).

### In Vivo Vascular Leakage Assay

Embryos from *Gars1*^ΔWHEP/+^ intercross were collected at E18.5 from timed pregnant females and then placed in petri dish. Each animal was placed directly on ice for 30-60 seconds, after which the anesthetized pup was injected in the temporal vein on the head under a dissecting microscope with 20 µL of 0.3% Evans blue solution[<u>29</u>]. Following 20 min incubation, residual Evans Blue dye within blood vessels was removed by intra-cardiac whole-body perfusion using physiological saline (pH 7.4). Lung was excised and put in 200 µL formamide for 48 hours at 60°C to extract the Evans blue dye. Afterwards, the amount of extravasated Evans Blue was quantified using a spectrophotometer at a wavelength of 620nm. Tissue samples were then dried for 96 hours at 60°C, weighed, and the amount of extravasated Evans Blue was normalized to its dry weight.

### Fluorescence-activated Cell Sorting (FACS)

To study G18’s effect on cell surface Nrp1, confluent MEFs seeded in 100 mm dishes were serum-starved for 4 hours. After wash by PBS twice, the cells were suspended in FACS buffer containing 5 mM EDTA in PBS with 1% BSA. Cells were treated with 16 μg/mL G18 for 0, 15, 30, and 60 minutes at 37°C with rotation. After centrifuge at 1200 rpm for 5 min, the supernatant was discarded. FACS buffer containing rat anti-mouse Nrp1 antibody conjugated with Alexa 647 (R&D, FAB599414, 1:100) or the control rat IgG conjugated with Alexa 647 (R&D, IC006R, 1:100) was used to resuspend cells and incubated in dark for 15 min. After washed twice in FACS buffer, the cells were finally resuspended in 100 μL FACS buffer and then analyzed by using NovoCyte 3000 with NovoSampler Pro instrument with BD FACS Diva 6 software and off-line analysis by using FlowJo software.

To detect cell surface VEGFR2 in MEFs, the same method was used except for no G18 treatment, and the antibodies used were rat IgG_2a_ kappa anti-VEGFR2 (CD309/FLK1) mAb, PE (ThermoFisher Scientific, #12-5821-82, FACS 1:100) with rat IgG_2a_ kappa isotype control, PE (ThermoFisher Scientific, #12-4321-80, FACS 1:100).

### Acidic Gel Northern Blot

Total RNA was extracted from E18.5 embryonic lung using TRIzol reagent (Invitrogen). For acetylated tRNA, the RNA pellet was dissolved in 10 mM sodium acetate (pH 4.5). For deacetylated tRNA, the RNA pellet was dissolved in 0.2 M Tris HCl (pH 9.5) and incubated for 1 hour at 37°C before precipitation with 75% ethanol and dissolving in 10 mM sodium acetate (pH 4.5). The above tRNA samples were loaded in home-made 6.5% urea gels in a big gel chamber (42 cm x 20 cm x 0.4 mm) and electrophoresed in 0.1 M sodium acetate (pH 5) as a running buffer overnight at 500 V in a cold room and then blotted onto Hybond-N+ membranes (GE Healthcare Life Sciences). The membrane was cross-linked under UV and then hybridized with 5’ biotinylated oligonucleotide probes at 65°C in hybridization buffer containing 20 mM sodium phosphate (pH 7.2), 300 mM NaCl, and 1% SDS. After washing in wash buffer containing 20 mM sodium phosphate pH 7.2, 300 mM NaCl, 0.1% SDS, and 2 mM EDTA, the membrane was incubated with HRP-conjugated streptavidin (Thermo Fisher) in hybridization buffer for 30 minutes. Chemiluminescent substrate was applied to the membrane, and tRNA signal was visualized in Biorad Chemidoc MP imaging system. The sequences of 5’ biotinylated oligonucleotide probes are as follows (from 5’ to 3’):

cytosolic tRNA^Gly-TCC^: TGCGTTGGCCGGGAATCGAACCCGGGTCAACTGCTTGGAAGGCAGCTATGCTCACCACTATACCACCAACGC-3’;

cytosolic tRNA^Gly-GCC^: TGCATKGGCCGGGAATCGAACCCGGGCCTCCCGCGTGGCAGGCGAGAATTCTACCACTGAACCACCMATGC;

mitochondrial tRNA^Gly^: TACTCTCTTCTGGGTTTATTCAGAATCTACTAATTGGAAGTCAGTTATATTAATTATACTAAG GGAGT.

### Confocal Microscopy

ECs were plated on glass coverslips coated with 1% gelatin from porcine skin (G9136, Sigma Aldrich) and allowed to adhere overnight to reach the required confluence to observe cell-to-cell contacts. Cells were washed in calcium (Ca^2+^) and magnesium (Mg^2+^) supplemented PBS, fixed for 10 minutes in PBS 2% paraformaldehyde (PFA), permeabilized in 0.1 % Triton X-100 for 2 min on ice. Cells were incubated with Goat polyclonal anti-VE-Cadherin (C-19 - Santa Cruz Biotechnology) primary Antibody diluted in PBS 1% donkey serum for 1h at room temperature and revealed by Alexa Fluor 555 donkey anti-goat IgG (H+L) secondary Ab (Invitrogen) diluted in PBS 1% donkey serum 1:100 for 30 minutes at room temperature. The slides were mounted on a microscope slide using Fluorescence Mounting Medium (DAKO) and were allowed to dry overnight at room temperature. Cells were analyzed by using a confocal laser-scanning microscope (TCS SP8 AOBS, Leica Microsystems, Mannheim, Germany). For image acquisition we used a 63X oil-immersion objective (N.A. 1.32). Image acquisition was performed by adopting the same laser power, gain, and offset settings for all the images of the same experiment and avoided saturation. We quantified the Mean Fluorescence Intensity of VE-cadherin localized in adherens junctions (AJs) drawing linear regions of interest (ROI) by means of the Leica Confocal Software Quantification Tool (Leica LAS-X).

### Statistical Analysis

Data are the mean +/-SEM of three biologically independent experiments. For statistical evaluation, parametric two-tailed Student’s *t* test was used to assess the statistical significance when two groups were compared, while parametric two-tailed analysis of variance (ANOVA) was applied when more than two groups were compared. Statistical differences were considered not significant (ns) = p-value > 0.05; significant * = p-value ≤ 0.05; ** = p-value ≤ 0.01; *** = p-value ≤ 0.001; **** = p-value ≤ 0.0001.

## Supporting Information

Supporting Information is available from the Wiley Online Library or from the author.

## Acknowledgements

The research leading to these results has received funding from: National Institutes of Health R35 GM139627 (to X.L.Y.), AIRC under IG 2023 - ID. 28763 and Ministero dell’Istruzione, dell’Università e della Ricerca (PRIN 2020EK82R5) (to G.S.). Y.T. was partially supported by a fellowship from National Foundation for Cancer Research.

## Author contributions

X.L.Y. conceived the project; X.L.Y., G.S., T.K., Y.T., and N.G. designed the experiments; X.L.Y., G.S., and T.K. supervised the research; H.Z., C.D.D., K.N.C., and J.Z. provided key reagents, methods, and technologies; Y.T., N.G., H.Z., M.H., K.N.C., and G.B. performed the experiments; Y.T., N.G., H.Z., M.H., S.K., M.H., K.N.C., J.Z., B.B., G.B., G.S., T.K., and X.L.Y analyzed the data; Y.T., N.G., S.K., M.H., K.N.C., B.B., G.B., G.S., T.K., and X.L.Y. interpreted the results; Y.T., N.G., B.B., G.S., T.K., and X.L.Y. wrote the paper; all authors read and approved the manuscript.

## Data Availability Statement

The data that support the findings of this study are available from the corresponding author upon reasonable request.

---

**Scheme 1.**
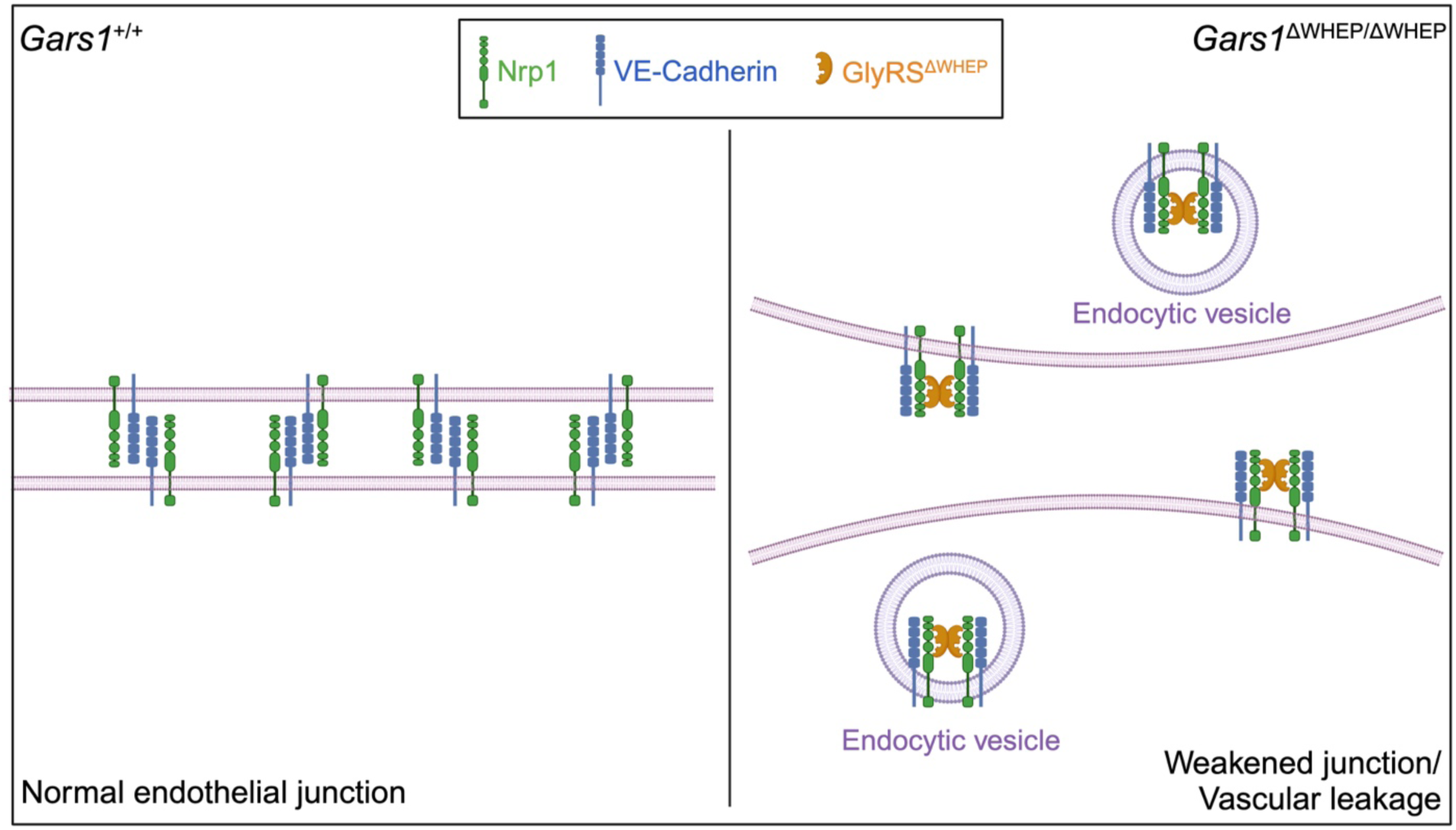
A schematic model explaining how vascular permeability is increased in *Gars1* ^ΔWHEP^ mice. Homophilic interactions between VE-cadherin molecules on adjacent endothelial cells are essential for maintaining endothelial cell-cell junction integrity. Neuropilin-1 (Nrp1) regulates the turnover of VE-cadherin at these junctions through its endocytic activity. A truncated form of glycyl-tRNA synthetase lacking the WHEP domain (GlyRS^ΔWHEP^) interacts with and activates Nrp1, promoting VE-cadherin internalization. This process disrupts endothelial cell-cell contacts and increases vascular permeability.

