## Supplemental figures for "WHEP Domain of Glycyl-tRNA Synthetase Regulates Neuropilin 1 Binding and Vascular Permeability"

SUPPLEMENTARY FIGURES AND FIGURE LEGENDS

Supplementary Figure 1

a

E12.5

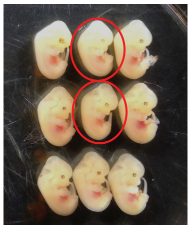

b

E12.5

| Genotype | Number of embryos | Observed ratio | Expected ratio |
|---|---|---|---|
| Gars1$^{\Delta WHEP/\Delta WHEP}$ | 10 | 23.8% | 25% |
| Gars1$^{\Delta WHEP/+}$ | 20 | 47.6% | 50% |
| Gars1$^{+/+}$ | 12 | 28.6% | 25% |
| Total | 42 | 100.0% | 100% |

c

E18.5

| Genotype | Number of embryos | Observed ratio | Expected ratio |
|---|---|---|---|
| Gars1$^{\Delta WHEP/\Delta WHEP}$ | 9 | 15.8% | 25% |
| Gars1$^{\Delta WHEP/+}$ | 34 | 59.6% | 50% |
| Gars1$^{+/+}$ | 14 | 24.6% | 25% |
| Total | 57 | 100.0% | 100% |

d

P0

| Genotype | Number of embryos | Observed ratio | Expected ratio |
|---|---|---|---|
| Gars1$^{\Delta WHEP/\Delta WHEP}$ | 10* | 14.1% | 25% |
| Gars1$^{\Delta WHEP/+}$ | 44 | 62.0% | 50% |
| Gars1$^{+/+}$ | 17 | 23.9% | 25% |
| Total | 71 | 100.0% | 100% |

*: 8 of the 10 pups were found dead at birth. 2 pups were alive at birth, but died soon after birth.

**Supplemental Figure S1. Phenotypes of homozygous *Gars1*$^{\Delta WHEP}$ mouse. a.** Representative images of mouse embryos at E12.5 from inter-crossing heterozygous *Gars1*$^{\Delta WHEP}$ mice. Red circles indicate homozygous *Gars1*$^{\Delta WHEP}$ embryos. **b-d.** Tables summarizing the number of embryos, the observed and expected ratio of wild type, heterozygous and homozygous *Gars1*$^{\Delta WHEP}$ mice at E12.5 (**b**), E18.5 (**c**), and P0 (**d**).

Supplementary Figure 2

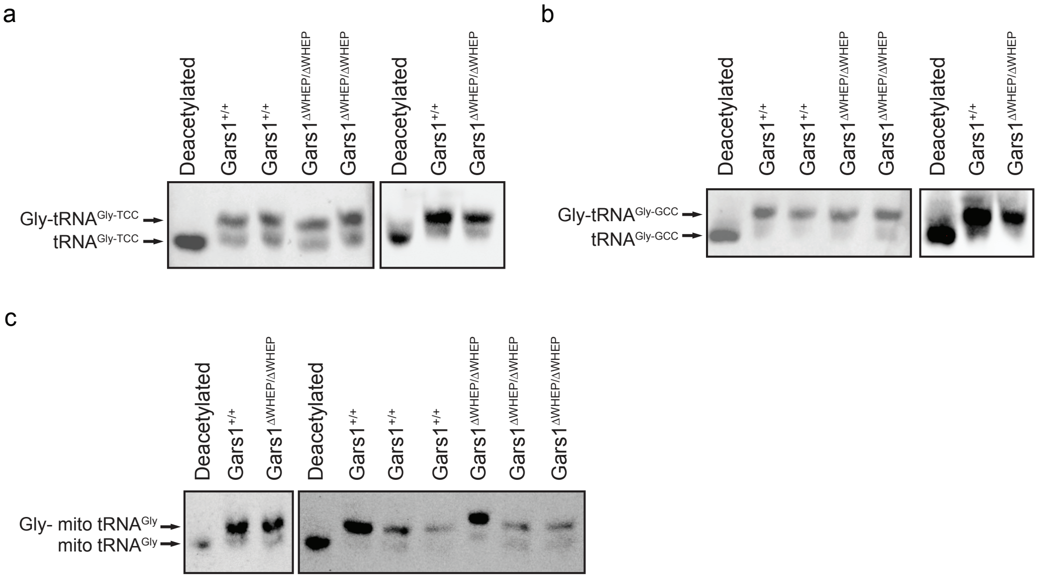


**Supplementary Figure S2. Acidic gel northern blot results to show in vivo charging status of cytosolic tRNA$^{Gly\text{-}TCC}$ (a), cytosolic tRNA$^{Gly\text{-}TCC}$ (b) and mitochondrial tRNA$^{Gly}$ (c) from lungs of E18.5 wild-type and *Gars1*$^{\Delta WHEP/\Delta WHEP}$ mouse embryos.**

Supplementary Figure 3

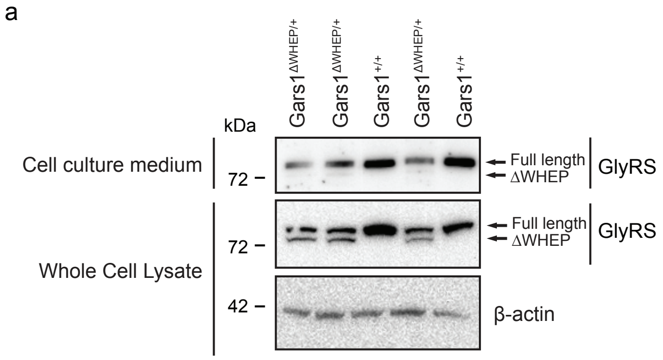


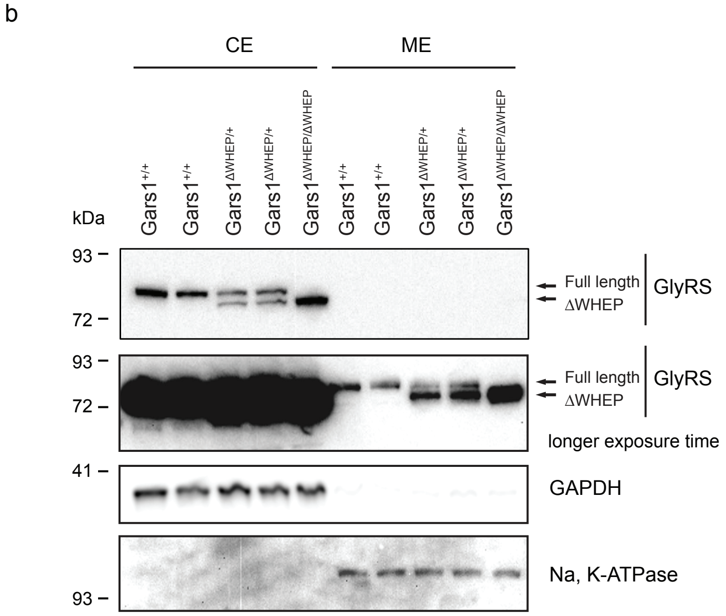


**Supplementary Figure S3. Secretion and cell fractionation analysis that include *Gars1$^{\Delta WHEP/+}$* MEFs. a.** GlyRS protein level from whole cell lysate (WCL) and culture medium of wild-type and *Gars1$^{\Delta WHEP/+}$* MEFs. **b.** Subcellular protein fractionation of wild-type, *Gars1$^{\Delta WHEP/+}$* and *Gars1$^{\Delta WHEP/\Delta WHEP}$* MEFs. CE: cytosol extraction, ME: membrane extraction. GAPDH and Na,K-ATPase were used as markers for CE and ME, respectively.

Supplementary Figure 4

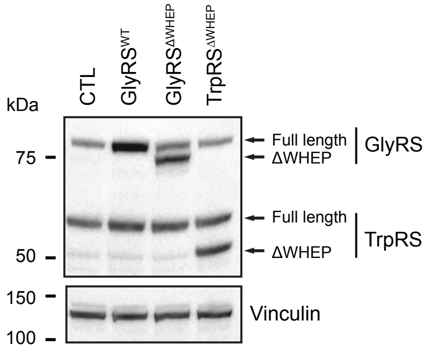


**Supplementary Figure S4. Overexpression of GlyRS$^{WT}$, GlyRS$^{\Delta WHEP}$ and TrpRS$^{\Delta WHEP}$ in primary cultures of human umbilical artery endothelial cells (HUAECs) through lentivirus transduction.** GlyRS$^{\Delta WHEP}$ indicates GlyRS$^{\Delta 19\text{-}52}$, missing 33 amino acids from the WHEP domain. TrpRS$^{\Delta WHEP}$ indicates TrpRS$^{\Delta 1\text{-}47}$, lacking the first 47 amino acids of the WHEP domain. Vinculin was used as internal control.

Supplementary Figure 5

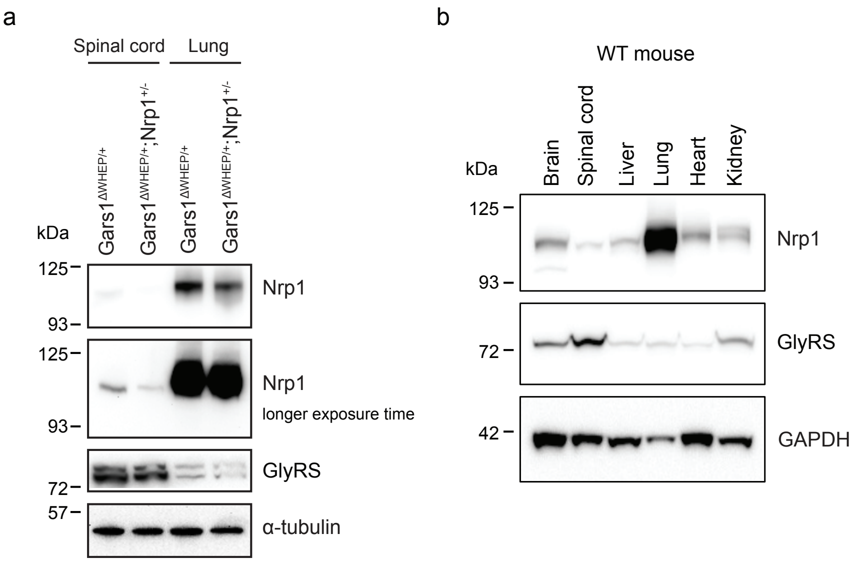


**Supplementary Figure S5. Nrp1 protein is enriched in the lung of mouse. a.** Western blot to show Nrp1 and GlyRS protein level in spinal cord and lung from adult *Gars1*$^{\Delta WHEP/+}$ and *Gars1*$^{\Delta WHEP/+}$;Nrp1$^{+/-}$ mouse, a-tubulin is used as internal control. **b.** Western blot to show Nrp1 and GlyRS protein level in different tissues from adult wild-type mice. GAPDH is used as internal control.

Supplementary Figure 6

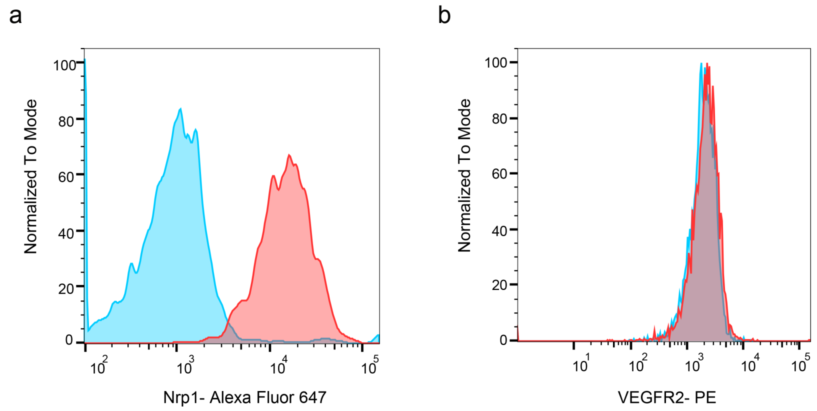


**Supplementary Figure S6. FACS analysis to show cell surface expression level of Nrp1 (a) and VEGFR2 (b) in MEFs.**

**Supplementary Video S1. Video shows that newborn *Gars1*$^{\Delta WHEP/\Delta WHEP}$ pup has difficulty to breathe.**
